# Expectations May Influence the Effects of Transcranial Direct Current Stimulation

**DOI:** 10.1101/279554

**Authors:** Sheida Rabipour, Allan D. Wu, Patrick S. R. Davidson, Marco Iacoboni

## Abstract

Growing interest surrounds transcranial direct current stimulation (tDCS) as a safe and inexpensive method for improving cognitive functions and mood. Nevertheless, tDCS studies rarely examine psychological factors such as expectations of outcomes, which may influence tDCS responsiveness through placebo-like effects. Here we sought to evaluate the potential influence of expectations on tDCS intervention outcomes. We assessed expectations of tDCS outcomes in 88 healthy young adults on three occasions: i) at baseline; ii) after reading information implying either high or low effectiveness of stimulation; and iii) after a single-session of sham-controlled anodal tDCS applied to the left dorsolateral prefrontal cortex, during working memory (WM) training. Participants were largely uncertain about the effectiveness of stimulation in improving cognitive function at baseline. High or low expectation priming using simple positive or cautionary messages significantly increased or decreased expectation ratings, respectively, but ratings significantly decreased following stimulation in all groups. We found greater improvement in participants who received high compared to low expectation priming. Participants who received active stimulation and low expectation priming exhibited the lowest performance, suggesting that expectation priming and stimulation may have interacted. We did not find a significant effect of baseline expectations, belief of group assignment, or individual characteristics on measures of WM and verbal fluency. However, controlling for baseline expectations revealed greater post-intervention improvement on the executive function measures in participants who received high (compared to low) expectation priming. People randomly assigned to receive high expectation priming reported having a more pleasant experience overall, including greater satisfaction. Our findings suggest that expectations of outcomes should be taken into account in tDCS-based experimental studies and clinical trials.

**Highlights:** - Based on prior knowledge, healthy subjects are uncertain about NIBS effectiveness.

- Expectations of NIBS can change after a single exposure to simple written messages.

- Expectations of outcomes may influence cognitive performance.

- High expectations may lead to a more positive experience and motivation to perform.

- Low expectations may be counterproductive to NIBS.

## Introduction

Non-invasive brain stimulation (NIBS) using magnetic fields or electrical currents is increasingly used in health and disease (Bennabi et al., 2014; Coffman, Clark, & Parasuraman, 2014; Kekic, Boysen, Campbell, & Schmidt, 2016). With low cost and relatively simple administration, NIBS holds understandable appeal for the investigation and modulation of cognitive mechanisms (Filmer, Dux, & Mattingley, 2014), treatment of clinical populations (Freitas et al., 2011; Gandiga, Hummel, & Cohen, 2006; Zhao et al., 2017) and specialized training of high-performing individuals (Scheldrup et al., 2014).

Transcranial direct current stimulation (tDCS) is a portable and generally safe (Bikson et al., 2016; Nitsche et al., 2008) form of NIBS alleged to enhance a vast set of functions. These include learning and implicit decision-making (Kincses et al., 2004), language production (Monti et al., 2013), episodic memory (Sandrini et al., 2014), and executive functions (Au et al., 2016; Dockery et al., 2009; McIntire, McKinley, Goodyear, & Nelson, 2014; Ruf, Fallgatter, & Plewnia, 2017).

Nevertheless, growing excitement among researchers, clinicians, and individual consumers is contrasted by a lack of reproducible findings across the tDCS literature (Berlim, Van den Eynde, & Daskalakis, 2013; Horvath, Forte, & Carter, 2015a, 2015b; Mancuso, Ilieva, Hamilton, & Farah, 2016; Medina & Cason, 2017; To, Hart, De Ridder, & Vanneste, 2016). In addition to considerable variability in reported effects, studies suggest high inter-individual differences in responsiveness to tDCS (Li, Uehara, & Hanakawa, 2015; Lopez-Alonso, Cheeran, Rio-Rodriguez, & Fernandez-del-Olmo, 2014). Factors such as electrode placement (Penolazzi, Pastore, & Mondini, 2013), strength of current (Chew, Ho, & Loo, 2015; Underwood, 2016), and stimulation schedule (Au et al., 2016) may contribute to differential effects, but do not appear to fully explain this variability. A recent study found that current intensities conventionally used in NIBS studies are likely insufficient to affect neuronal circuits directly, suggesting that reported behavioural and cognitive effects may result from indirect mechanisms (Voroslakos et al., 2018). Moreover, some researchers have argued that published tDCS studies for cognition and working memory (WM) are underpowered and have little evidential value (Medina & Cason, 2017). These findings highlight the uncertainty about the mechanisms through which tDCS influences cognition and behaviour (Farah, 2015; Walsh, 2013), as well as the need for greater scientific rigour in evaluations of tDCS (Riggall et al., 2015).

Crucially, studies of tDCS have rarely examined psychological factors such as expectations of outcomes, which may influence tDCS responsiveness through placebo or Hawthorne-like effects (McCambridge, Witton, & Elbourne, 2014; Shiozawa, Duailibi, da Silva, & Cordeiro, 2014). Evidence for the influence of expectations on cognitive interventions (Foroughi, Monfort, Paczynski, McKnight, & Greenwood, 2016; Rabipour, Andringa, Boot, & Davidson, 2017; Rabipour & Davidson, 2015) and performance (Schwarz, Pfister, & Buchel, 2016), coupled with the possible influence of factors such as emotional state (Sarkar, Dowker, & Cohen Kadosh, 2014) and motivation (Jones, Gozenman, & Berryhill, 2015) on responsiveness to tDCS, underline the importance of examining the potential influence of expectations on tDCS. Furthermore, such expectations may at least partially account for the individual differences in tDCS responsiveness as well as variability in reported tDCS effects.

In the present study we assessed expectations of NIBS and examined the effects of expectation priming on cognitive performance following tDCS in healthy young adults (see also Rabipour et al., 2017; Rabipour & Davidson, 2015). Specifically, we asked: i) whether people tend to have neutral, optimistic, or pessimistic expectations of NIBS; ii) whether expectations of NIBS outcomes can be modified based on information indicating that the procedure either can or cannot enhance cognitive function; and iii) whether expectations of NIBS interact with the effects of anodal tDCS during performance of a cognitively challenging task. Our study represents a conceptual replication and extension of previous research reporting enhanced WM performance following a single session of anodal tDCS applied to the left dlLFC, and showing greater within- and between-session improvements following concurrent (i.e., “online”) rather than sequential (i.e., “offline”) task performance (Martin, Liu, Alonzo, Green, & Loo, 2014; see also Ruf et al., 2017).

## Materials and Methods

### Participants

We recruited 90 healthy young adults with normal or corrected-to-normal vision and hearing from the Psychology subject pool at the University of California, Los Angeles (UCLA). Participants were enrolled during an initial single-blind round of data collection: *n* = 52, 31 women, age=20.47±1.85 years, with one woman excluded following in-person screening; and during a follow-up double-blind round: *n* = 38, 22 women, age=20.61±3.35 years, with one woman excluded due to technical difficulties preventing completion of testing. We screened participants based on the following exclusion criteria: i) seizure disorder; ii) history of serious head trauma; iii) presence of metal, other than dental work, in the head; iv) presence of foreign object in body (e.g., pacemaker); v) electroconvulsive therapy in the past six months; vi) current pregnancy; vii) hospitalization for psychiatric reasons in the past six months; viii) change in antipsychotic medication in the past three months; ix) a major depressive episode in the past month; or x) active abuse of drugs or alcohol in the past six months. We received ethical approval to conduct this study from the UCLA Institutional Review Board and the University of Ottawa Research Ethics Board.

### Experimental Protocol

After providing consent, participants were assigned to one of two expectation priming conditions: i) High expectation priming, in which participants were told they would receive a type of brain stimulation known to improve performance (Fig. 1A); and ii) Low expectation priming, in which participants were told they would receive a type of brain stimulation with no known benefits (Fig. 1B). We then randomized participants to receive one of two stimulation conditions: active anodal or sham tDCS. All participants proceeded to complete a single session of tDCS based on their randomized group assignment (Table 1) while performing an online WM task.

**Table 1.** Random assignment of participants (*n* women) into the experimental conditions based on recruitment as part of an initial single blind or follow-up double blind protocol.

|  | **Active, High** | **Active, Low** | **Sham, High** | **Sham, Low** |
| --- | --- | --- | --- | --- |
| Single blind | 12 (8) | 12 (7) | 14 (8) | 13 (8) |
| Double blind | 12 (4) | 9 (8) | 6 (4) | 10 (5) |

**Fig. 1.**
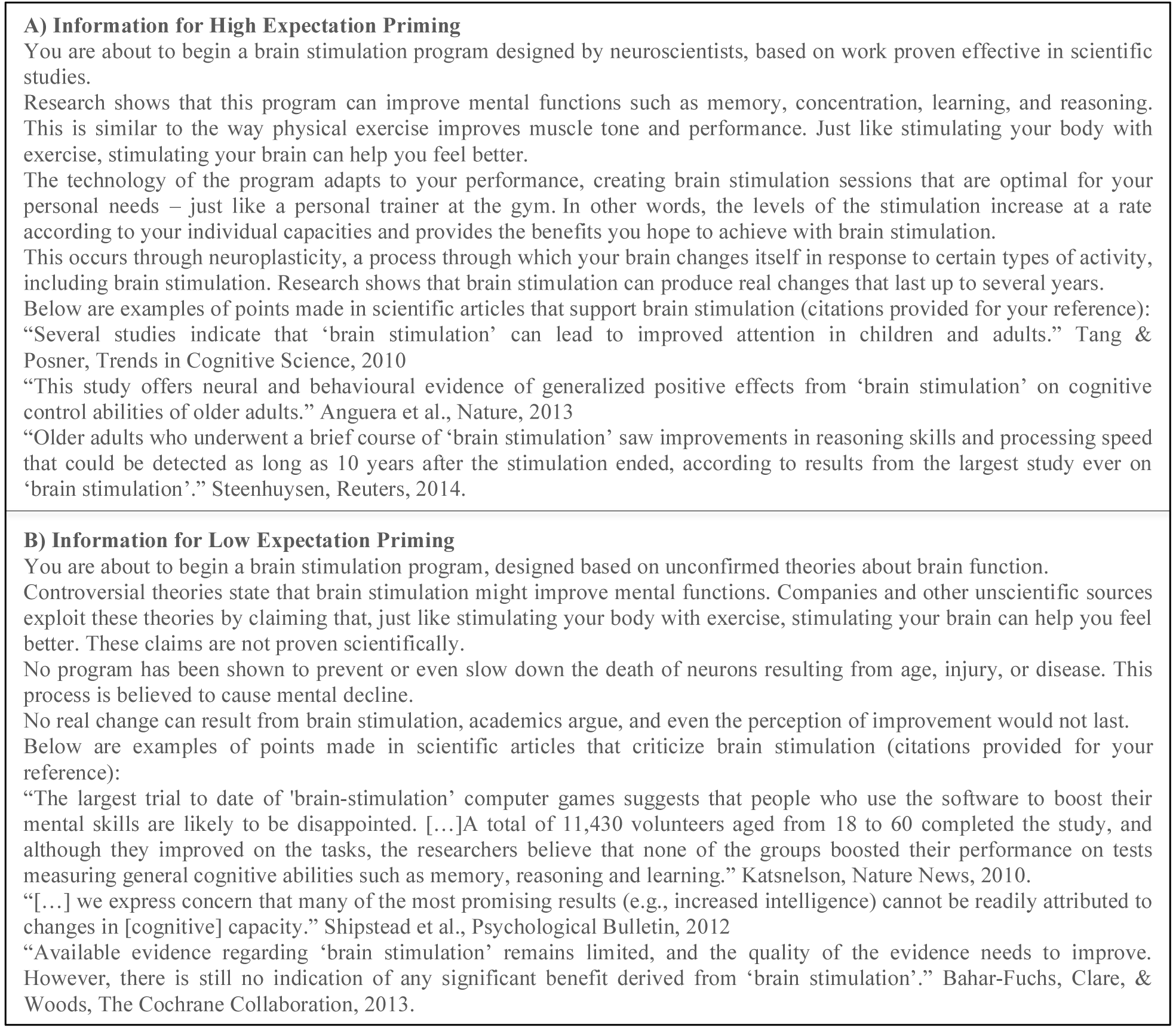
A) High and B) Low expectation priming messages delivered to participants.

### Expectation Assessment

Participants rated their expectations of tDCS effectiveness using the Expectation Assessment Scale (EAS), described in our previous work (Rabipour et al., 2017; Rabipour & Davidson, 2015; Rabipour, Davidson, & Kristjansson, 2018), on three occasions: i) at baseline; ii) after receiving High or Low expectation priming; and iii) after the tDCS session. Briefly, participants rated their expectations of outcomes on a scale from 1-7 (1 = “completely unsuccessful,” 2 = “fairly unsuccessful,” 3 = “somewhat unsuccessful,” 4 = neutral/“I have absolutely no expectations,” 5 = “somewhat successful,” 6 = “fairly successful,” 7 = “completely successful”). We probed expectations on nine outcomes: i) general cognitive function; ii) memory; iii) concentration; iv) distractibility; v) reasoning ability; vi) multitasking ability; vii) performance in everyday activities; viii) task accuracy; and ix) response time. Due to missing data for “concentration”, “task accuracy”, and “response time”, we excluded those domains from our repeated-measures analyses.

In the event of participants probing further about the study objectives and hypotheses, researchers were instructed to respond in line with the nature of the participant’s assigned expectation priming condition: for participants who received high expectation priming, experimenters explained that NIBS has been shown to be effective and the study aims to confirm previously reported effects; for participants who received low expectation priming, researchers explained that the body of research is contentious and that the present study aims to confirm no effect of the intervention. During data collection, however, participants rarely inquired further about the study after reading the expectation priming messages.

### Stimulation Procedure

During our initial single-blind round of data collection, we delivered tDCS through a battery-driven constant direct current stimulator (ActivaDose II, ActivaTek Inc, USA) using a pair of electrodes in 5 x 7cm saline-soaked synthetic sponges. The anodal electrode was placed at the F3 scalp location based on the International 10/20 system. The cathode was placed along the supraorbital nerve, 2-3cm lateral to the nasion-inion line on the contralateral side. The electrodes were held against the head with a standard head strap provided by Soterix Medical and an additional custom strap of Lycra synthetic fibre. For both active and sham conditions, the current was turned on and increased in a ramp-like fashion over approximately 30 seconds until reaching a strength of 2.0mA, with a current density equal to 0.04mA/cm^2^. Stimulation in the active tDCS condition was maintained for 20 minutes at 2.0mA, and then ramped down to 0.0mA over 30 seconds. For participants in the sham condition, the current was applied for only 30s and then ramped down to 0.0mA over 30 seconds. This allowed participants in each group to experience the same initial sensations (i.e., mild tingling). Participants habituate rapidly to the tingling sensations on the scalp, thus preserving the sham manipulation (Poreisz, Boros, Antal, & Paulus, 2007). The safety and potential cognitive effects of these stimulation parameters have been previously validated in healthy participants (Bikson et al., 2016; Iyer et al., 2005).

We recruited additional participants to undergo the same procedure using a NeuroConn Direct-Current Stimulator Plus (neuroCare Group, GmbH) with pre-programmed codes to deliver active vs. sham conditions in a double-blind manner. Stimulation montage and ramping conditions were identical in both phases, with the exception that current was held at 2.0mA for 40s in the sham condition before ramping down to 0.0mA.

### Online Working Memory Task

All participants performed a custom designed n-back task for 20 minutes, concurrent with active or sham tDCS. In our initial round of data collection, participants performed three different types of n-back, presented randomly over the course of the 20 minutes. Specifically, the task required participants to identify either: i) a letter or number repeated after n = 3 frames, among a random sequence of letters and numbers; ii) an image repeated after n = 3 frames, among a random series of images; or iii) the spatial position of a single image moving across a 3 x 3 grid, repeated after n = 3 frames.

The n-back exercises were originally designed for a five-week, multi-session cognitive training program. Each type of n-back task was subdivided into trial blocks lasting 180s (letter-number and spatial type of n-back; 150 trials within each block) or 1080s (image type of n-back; 900 trials within each block) in total, wherein stimuli were presented in immediate succession and participants had 1200ms to respond. For the majority of participants (*n* = 64), the software randomly generated the type of n-back task that each participant completed. Technical issues with the software prevented us from accessing data pertaining to the nature of correct responses (i.e., hits vs. correct rejections). After updating the database to rectify these issues, we removed the randomization of n-back types and instructed the remaining participants (*n* = 24) to complete only the letter-number n-back task.

During both rounds of data collection, participants were first allowed to practice with a 2- back task and then graduated to a 3-back task, until comfortable with the rules. Practice time was maintained within 10 minutes. The experimenter manually terminated the task once the stimulation protocol was complete. Due to technical difficulties with the software, we were unable to store complete data for 16 participants (single blind: *n* = 1; double blind: *n* = 15).

### Neuropsychological Assessment

To assess changes in cognitive performance, we administered a brief battery of neuropsychological tests measuring executive functioning, with a focus on working memory and verbal fluency, both before and after receiving brain stimulation. All of the administered tests were timed with a digital stopwatch.

**Letter-Number Sequencing Task:** The letter-number sequencing (LNS) task, a subtest of the Wechsler Adult Intelligence Scale, is a brief, standardized measure of executive function that indexes verbal and visuospatial WM performance, as well as processing speed (Crowe, 2000). During the task, examiners read a series of scrambled letters and numbers to participants, who must then recite the numbers in sequential order, followed by the letters in alphabetical order. **Digit Span Task:** The digit span task, a simple span subtest of the Wechsler Adult Intelligence Scale, includes a forward and backward digit span task. For the forward digit span (FDS) task, a measure of short-term recall, participants repeated a string of numbers that the examiner read aloud; in the backward digit span (BDS) task, an index of WM capacity and incomplete effort (Axelrod, Fichtenberg, Millis, & Wertheimer, 2006), participants recited the string of numbers in the reverse order.

**Controlled Oral-Word Association Test (COWAT) and Animal Naming Test:** The COWAT measures letter fluency (LF) or knowledge (i.e., letter-sound associations) and requires the generation of words that begin with a given letter within 60 seconds (Lanting, Haugrud, & Crossley, 2009). In the present study, participants were asked to name as many words as possible that begin with the letter “C”, then “F”, and finally “L”, or “P”, then “R”, and finally “W” at baseline and following tDCS, in counterbalanced order. The animal-naming test, administered immediately after the COWAT, is a measure of semantic fluency (SF) that requires participants to name as many animals as possible in 60 seconds. In the present study, scores represent the total number of admissible words reported.

### Feedback on Perceived Experience

Immediately following the stimulation procedure, we asked participants to report on their experience. On a scale from 1-7 (1 = “very strongly disagree,” 2 = “strongly disagree,” 3 = “disagree,” 4 = neutral/“neither agree nor disagree,” 5 = “agree,” 6 = “strongly agree,” 7 = “very strongly agree”), participants rated the degree to which they found the experience: i) enjoyable; ii) challenging; iii) frustrating; iv) engaging; v) boring; vi) motivating; and vii) satisfying.

### Statistical Analysis

Using G*Power version 3.1, we determined that a total sample of 84 participants would detect a moderate effect size at an alpha level of .05 with 80% power.

Primary outcomes were analysed using repeated-measures analysis of variance (ANOVA), as well as univariate ANOVA and multivariate ANOVA (MANOVA), with within-between factors interactions to evaluate the outcome measures described, at an alpha level of 0.05. For our online WM task, we analysed the proportion of correct trials (i.e., hits and correct rejections) and of false alarms (i.e., incorrect clicks as a proportion of total incorrect trials). For our transfer measures, we combined the behavioural outcome measures a priori into a single standardized “executive function” score at baseline and following stimulation. We used the mean and standard deviation from baseline scores to compute the composite for both baseline and post-stimulation scores. As a more conservative approach, we initially opted to analyse our results with condensed composite scores comprised of only the LNS, BDS, and LF total scores, which have been shown to correlate more strongly with executive function (e.g., see: Crowe, 2000; McCabe, Roediger, McDaniel, Balota, & Hambrick, 2010; Redick & Lindsey, 2013; St Clair-Thompson, 2010). Based on feedback from a reviewer, we report results from both analyses (i.e., with the complete composite scores and the condensed composite scores). We employed Pearson correlations and Principal Components Analysis (PCA) of baseline scores to verify the relatedness of these outcomes in our sample. Where applicable, we used the Holm-Bonferroni correction for multiple comparisons and Greenhouse-Geisser correction for sphericity. We performed analyses using IBM SPSS Statistics, Inc. version 23 and R version 3.1.1.

## Results

Expectation ratings and measures of performance did not significantly differ based on blinding (i.e., single vs. double). We therefore report results from both recruitment phases together.

### How do people perceive non-invasive brain stimulation?

Based on their prior knowledge and perceptions of NIBS, participants were largely uncertain of the effectiveness of NIBS (Fig. 2). Responses were significantly above neutral only for “memory” in participants assigned to receive low expectation priming and sham tDCS (*M* = 4.57, *SD* = 0.788, t_[22]_ = 3.44, *p* = 0.002, Cohen’s *d* = 1.47; Fig. 2d) after correcting for multiple comparisons in a series of single-sample t-tests comparing each of the baseline ratings to the neutral point (rating of “4”). Because participants had not yet received the priming messages, this significant group difference in the baseline expectation ratings may represent a chance finding unrelated to our experimental protocol.

**Fig. 2.**
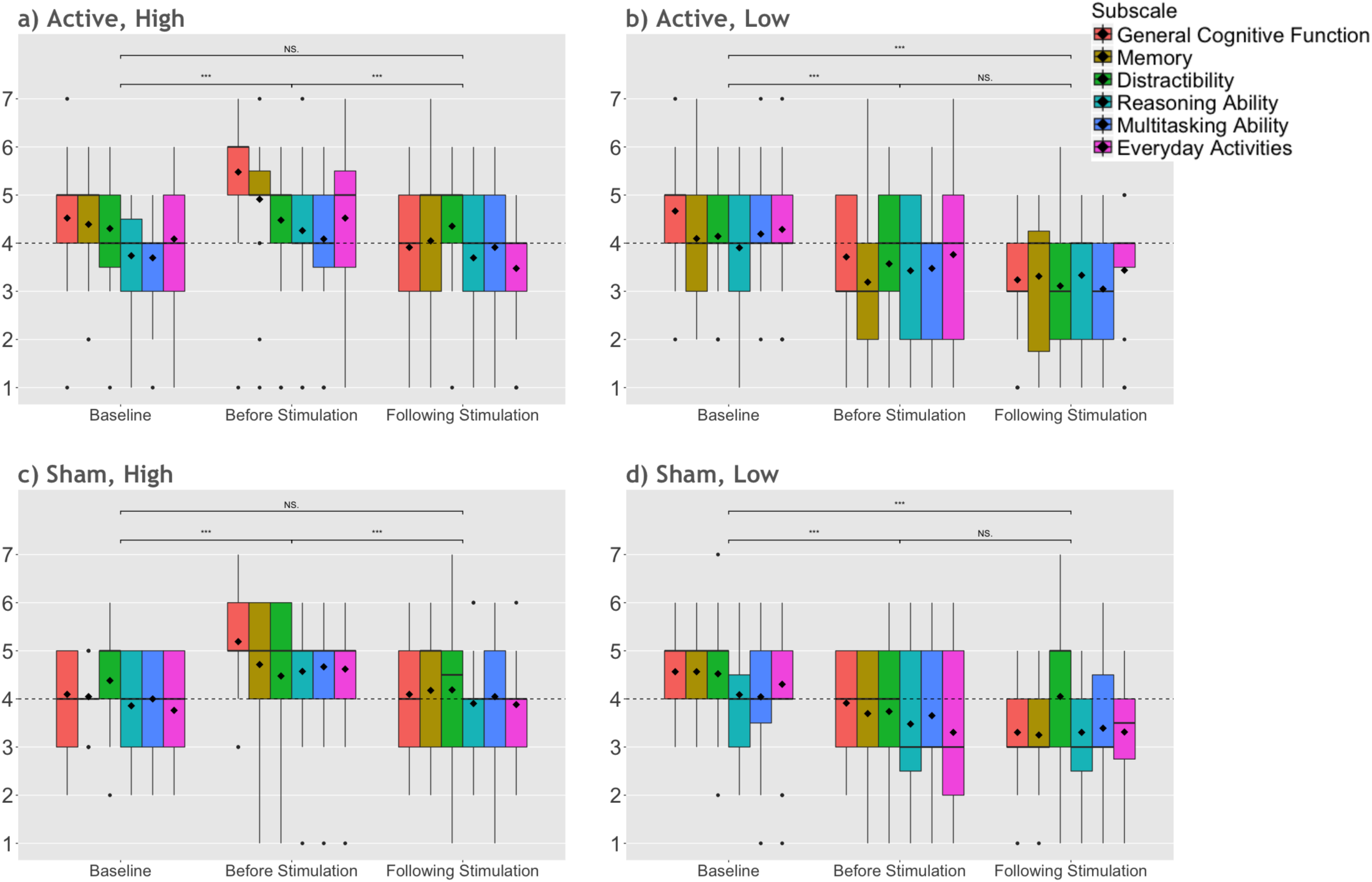
Mean group ratings of expected tDCS effectiveness for improving performance in participants who received a) active stimulation and high expectation priming, b) active stimulation and low expectation priming, c) sham stimulation and high expectation priming, and d) sham stimulation and low expectation priming. The upper and lower whiskers represent 1.5 x the inter-quartile range. Dashed lines indicate a neutral score (rating of “4”); bold lines indicate group medians; diamonds indicate group means. \**p* < .05. \*\**p* < .01. \*\*\**p* < .001.

We performed repeated measures ANOVA examining expectation ratings on each of the cognitive domains (“general cognitive function”, “memory”, “distractibility”, “reasoning ability”, “multitasking ability”, “performance in everyday activities”) across time (baseline, before simulation, and following stimulation) based on priming (high vs. low) and stimulation condition (active vs. sham). This analysis revealed a significant effect of time (*F*_[1.5,73.66]_ = 7.81, *p* = .002, η_p_^2^ = .137), of our priming conditions (*F*_[1,49]_ = 10.562, *p* = .002, η_p_^2^ = .177) and of cognitive domain (*F*_[4.09,200.2]_ = 3.745, *p* = .005, η_p_^2^ = .071) on expectation ratings, a significant interaction between time and priming condition (*F*_[2, 73.66]_ = 10.147, *p* < .0001, η_p_^2^ = .172), and a three-way interaction between time, cognitive domain, and priming condition (*F*_[10,490]_ = 1.997, *p* = .032, η_p_^2^ = .039; Fig. 2).

Follow-up repeated measures ANOVA comparing ratings based on each priming condition across time revealed a significant effect of time (*F*_[1.4,35]_ = 7.022, *p* = .002, η_p_^2^ = .219) and of domain (*F*_[3.6,91.19]_ = 4.566, *p* = .003, η_p_^2^ = .154) as well as an interaction between time and domain (*F*_[10,250]_ = 2.108, *p* = .024, η_p_^2^ = .078) in participants who received high expectation priming. In participants who received low expectation priming, we found only a significant effect of time (*F*_[1.6,38.16]_ = 10.917, *p* < .0001, η_p_^2^ = .313). Overall, our pattern of results suggests that expectations increased before stimulation (i.e., after receiving the priming information) and decreased following stimulation in participants who received high expectation priming; conversely, in participants who received low expectation priming, expectations decreased relative to baseline before stimulation and remained low after stimulation (see Supplementary Table S1 for expectation ratings across domain, within each group).

### Does expectation priming interact with stimulation effects on cognitive performance?

#### Online Working Memory Task

Participants in all groups completed approximately four trial blocks in total during stimulation (*M* = 4.11, *SD* = 1.68; *ns*). For sessions where trial blocks were randomized across the three types of n-back, the number of trial blocks completed in each type of n-back did not differ between groups (Table 2a).

**Table 2.** a) Mean number of n-back trial blocks completed during stimulation and b) proportion of correct responses, based on n-back type, for participants who received all three types of n-back.

| a) | Letter-Number n-back | Image n-back | Spatial n-back |
| --- | --- | --- | --- |
| Active, High | 1.27 ± 1.44 | .26 ± .46 | 1.07 ± 1.39 |
| Active, Low | 1.29 ± 1.44 | .36 ± .50 | .93 ± .92 |
| Sham, High | 1.56 ± 1.50 | .19 ± .40 | 1.31 ± 1.25 |
| Sham, Low | 1.61 ± 1.72 | .33 ± .49 | .78 ± 1.00 |
| b) |  |  |  |
| Active, High | .92 ± .07 | .96 ± .08 | .95 ± .06 |
| Active, Low | .89 ± .09 | .92 ± .06 | .91 ± .07 |
| Sham, High | .87 ± .10 | .97 ± .40 | .89 ± .09 |
| Sham, Low | .92 ± .08 | .94 ± .49 | .93 ± .08 |

ANOVA examining n-back performance^1^ across expectation priming (high vs. low priming) and stimulation condition (active vs. sham) revealed a significant interaction between expectation and stimulation on correct trials (i.e., hits and correct rejections; *F*_[1, 68]_ = 9.002, *p* = .004, η_p_^2^ = .117; Fig. 4). Specifically, participants who received low expectation priming and active tDCS had significantly fewer correct trials (i.e., correct identification of targets and correct rejection of distractors) compared to their counterparts who received high expectation priming and active tDCS (*t*_(20.28)_ = 2.754, *p* = .012; *M*_*diff*_ = .085, *CI* = .02-.149; Cohen’s *d* = 1; Fig. 3a).

**Fig. 3.**
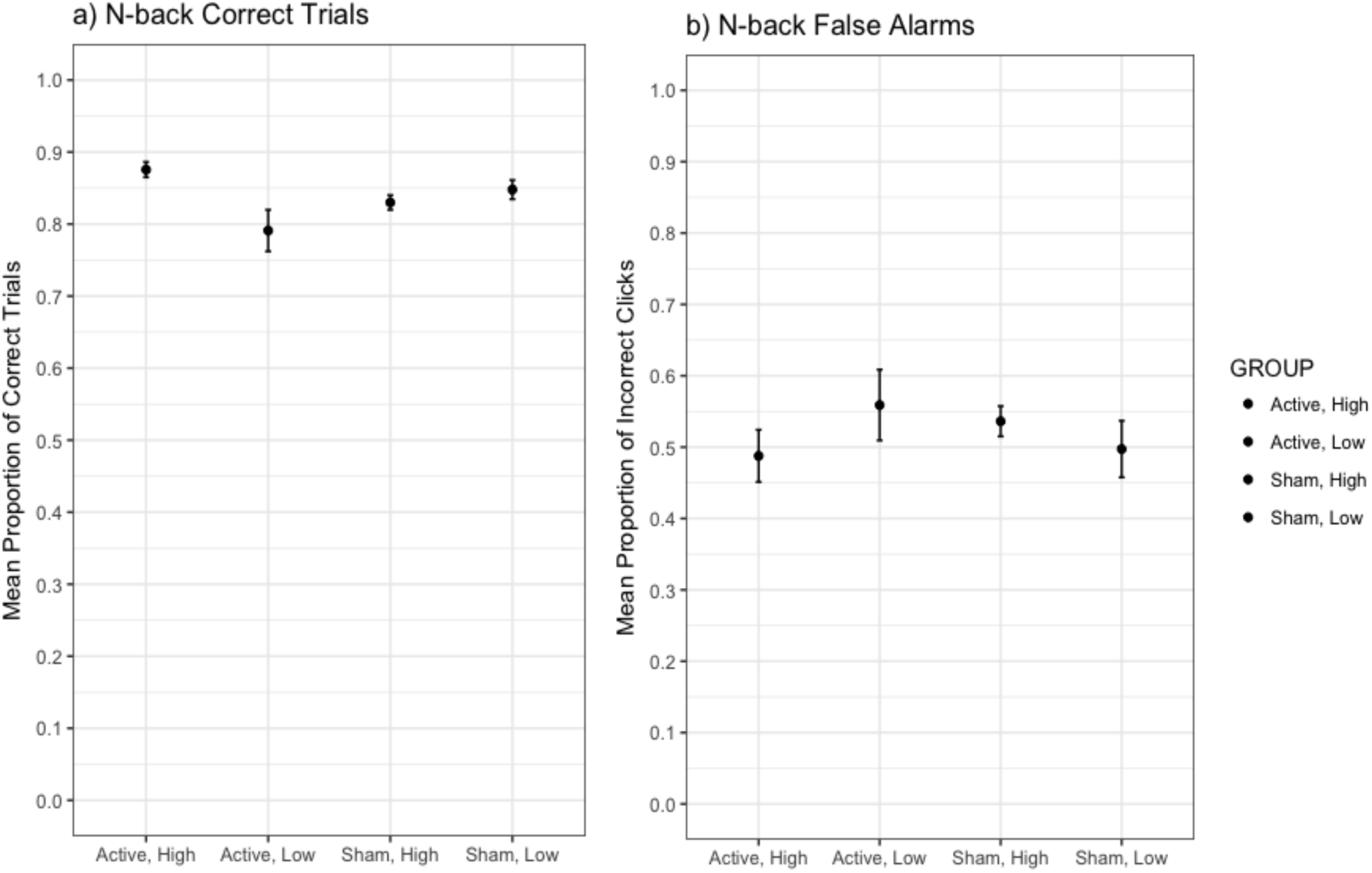
Performance on n-back task represented as a) correct trials (i.e., correct identification of targets and rejection of distractors) and b) false alarm rate (i.e., incorrect identification of distractors as targets, as a proportion of incorrect trials). Results indicate that, among participants who received active tDCS, the proportion of correct trials was significantly higher in participants who received high expectation priming compared to those who received low expectation priming. Despite greater variability in scores of participants who received active stimulation and low expectation priming, these results are not driven by outliers. Error bars represent standard error of the mean. Due to technical issues with task programming, we were unable to separate hits and correct rejections.

**Fig. 4.**
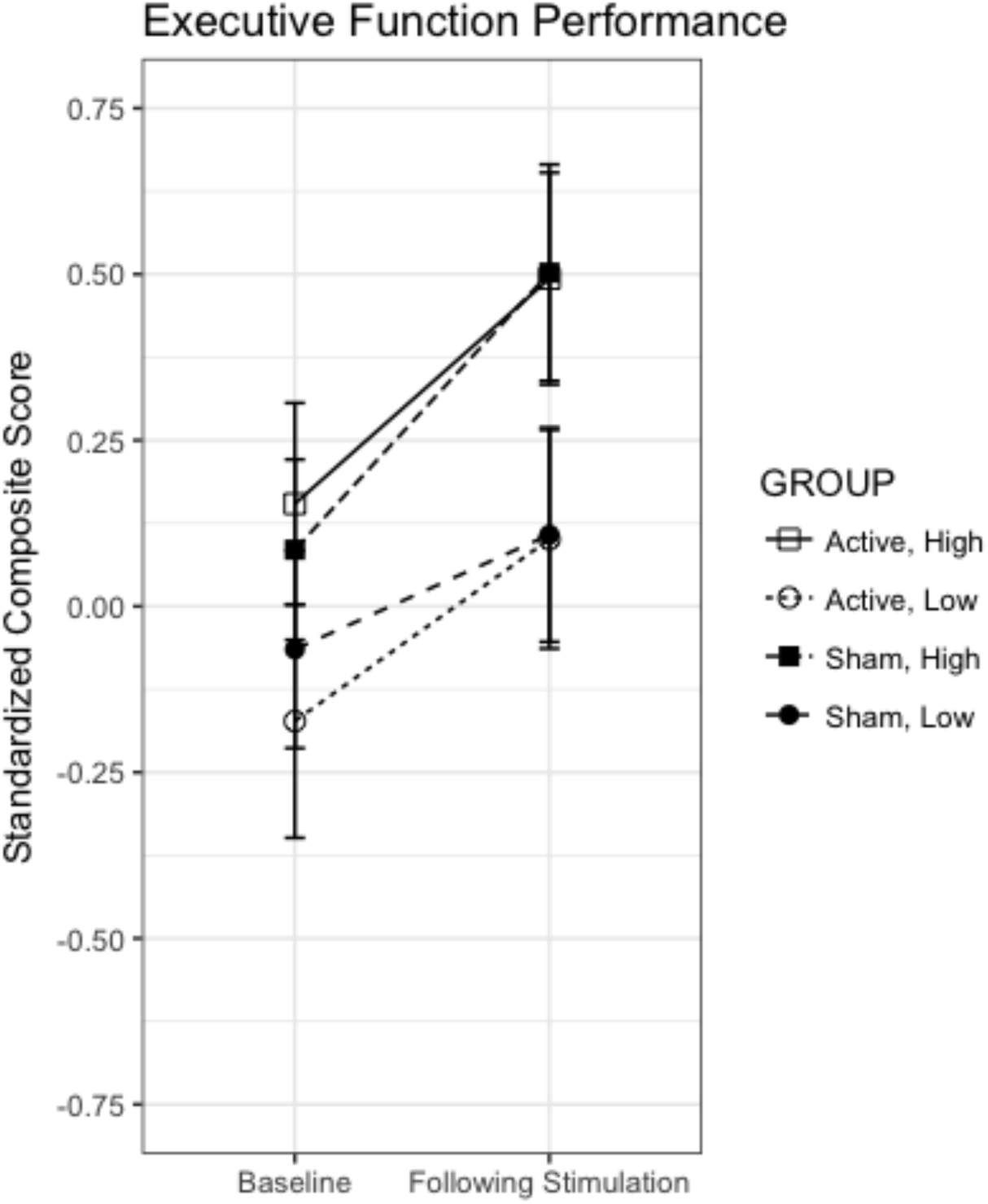
Executive function performance represented as standardized composite scores of all transfer tests at baseline and following stimulation. Results indicate a significant interaction between time, expectation priming, and stimulation condition. Bars represent standard error of the mean.

**Fig. 5.**
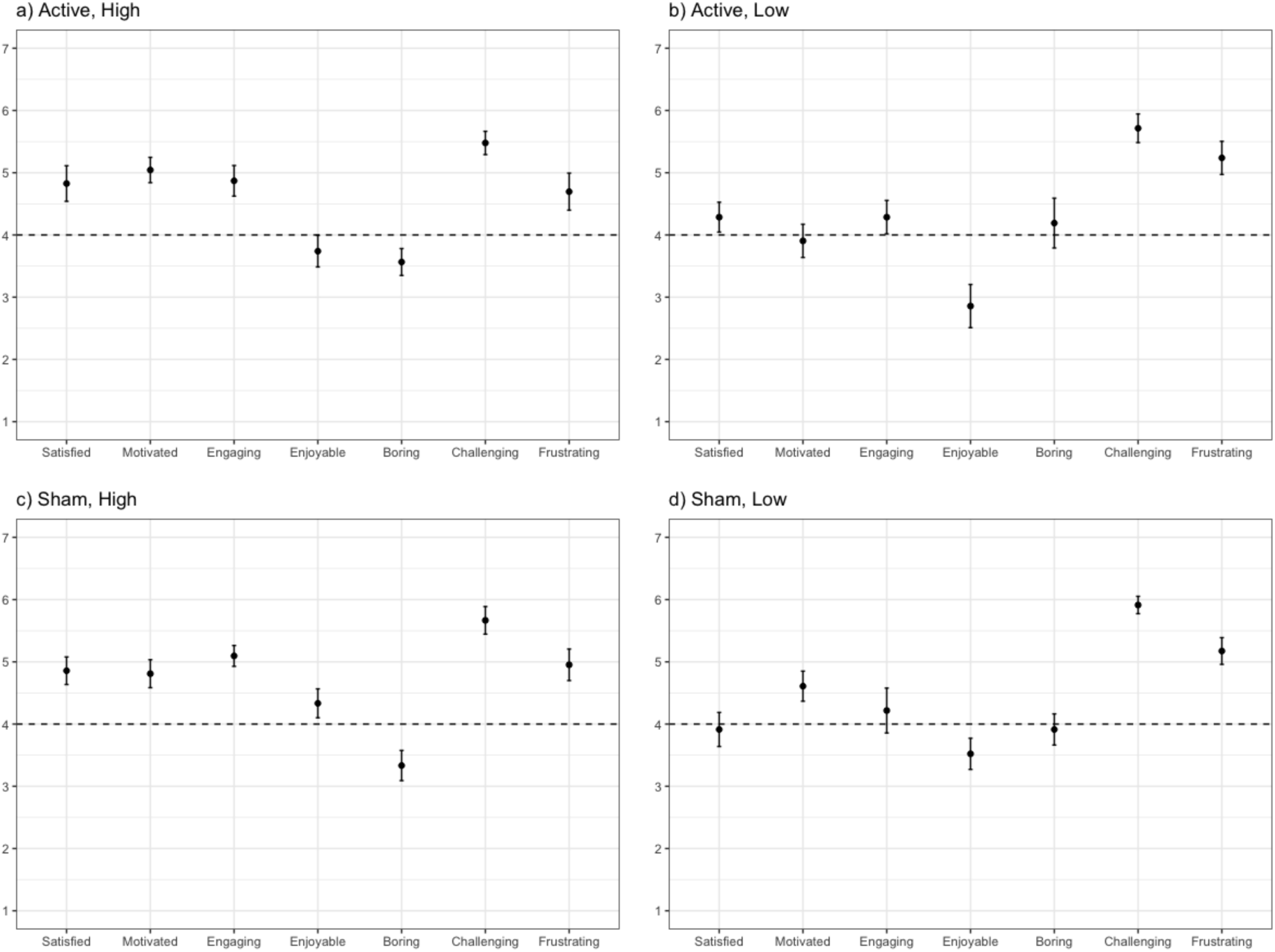
Feedback on perceived experience.

These differences remained significant after accounting for baseline expectations and performance on neuropsychological tests, as well as anticipated and experienced side effects. We found a comparable, albeit non-significant, pattern for false alarms (i.e., incorrect identification of distractors as targets as a proportion of incorrect trials; Fig. 3b). However, we found no main effect of expectation priming (*F*_[1, 68]_ = 3.819, *p* = .055) or stimulation condition (*F*_[1, 68]_ = .107, *p* = .744) on n-back performance.

For participants who completed all three types of n-back, we included n-back type as a within-subjects variable in a separate ANOVA of performance, as measured by correct responses. We found a significant main effect of n-back type (*F*_[1.63, 96.04]_ = 3.752, *p* = .035, η_p_^2^ = .06), but no interaction between n-back type and expectation (*F*_[1.63, 96.04]_ = 1.065, *p* = .348) or stimulation condition (*F*_[1.63, 96.04]_ = .679, *p* = .48), and no three-way interaction between these variable (*F*_[1.63, 96.04]_ = .008, *p* = .697). Following up on the effect of n-back type revealed that performance was significantly higher on the image n-back compared to the letter-number n-back (*t*_(63)_ = 2.38, *p* = .02; *M*_*diff*_ = .046, *CI* = .007-.085; Cohen’s *d* = .60; Table 2b). As in our previous analysis, we also found a significant interaction between expectation priming and stimulation condition (*F*_[1,59]_ = 7.38, *p* = .009, η_p_^2^ = .11), wherein participants who received low expectation priming and active tDCS had significantly fewer correct trials compared to those who received high expectation priming and active tDCS (*t*_(27)_ = 2.553, *p* = .017; *M*_*diff*_ = .058, *CI* = .023-.105; Cohen’s *d* = .98).

#### Executive Function Transfer Tasks

Pearson correlations demonstrated significant positive correlations between most executive function scores (r ≥ .261, *p* < .014), with the exception of BDS and SF (r=.192, *ns*). PCA indicated that all scores loaded onto a single component, explaining 53.56% of the sample variance.

Repeated measures ANOVA of priming (high vs. low) and stimulation (active vs. sham) condition across time (i.e., before and after the stimulation procedure) revealed a significant effect of time (*F*_[1, 84]_ = 64.12, *p* < .0001, η_p_^2^ = .433), of expectation priming (*F*_[1,84]_ = 4.169, *p* = .044, η_p_^2^ = .047), and an interaction between time and expectation priming (*F*_[1, 84]_ = 4.468, *p* = 0.037, η_p_^2^ = .051) on the standardized executive function composite score (Fig. 4). Closer examination of the interaction between time and expectation priming revealed greater improvement in participants who received high expectation priming, compared to those assigned to low expectation priming; although performance was not significantly different between the two priming conditions at baseline (*t*_(86)_ = 1.559, *p* = .123; *M*_*diff*_ *=* .237, *CI* = .065-.54), participants in the high expectation condition performed better than those in the low expectation condition following stimulation (*t*_(86)_ = 2.459, *p* = .016; *M*_*diff*_ *=* .394, *CI* = .076-.713; Cohen’s *d* = .53). We did not find a main effect of stimulation (*F*_[1,84]_ = .001, *p* = .973), interactions between time and stimulation (*F*_[1,84]_ = .001, *p* = .977) or expectation priming and stimulation (*F*_[1,84]_ = .167, *p* = .684), or a three-way interaction between time, priming, and stimulation (*F*_[1,84]_ = 1.975, *p* = .164).

We re-analyzed these results in an ANCOVA, with the inclusion of baseline ratings on all 6 cognitive domains as covariates. We found the same general pattern of results, with the following differences: a smaller effect of time (*F*_[1, 78]_ = 6.24, *p* = .015, η_p_^2^ = .074) and marginal effect of priming (*F*_[1, 78]_ = 6.24, *p* = .06), but larger interaction between time and priming (*F*_[1, 78]_ = 5.27, *p* = .024, η_p_^2^ = .063).

Performing the ANOVA on the condensed executive function composite score (i.e., including only the LNS, BDS, and LF scores) demonstrated a significant effect of time (*F*_[1, 84]_ = 46.345, *p* < .0001, η_p_^2^ = .355) and a subtle interaction between time and expectation priming (*F*_[1,_ 84] = 3.905, *p* = 0.051, η_p_^2^ = .044). Closer examination of the interaction revealed greater improvement in participants who received high expectation priming, compared to those assigned to low expectation priming. Specifically, although baseline performance did not significantly differ between the two priming conditions (*t*_(86)_ = 1.231, *p* = .222; *M*_*diff*_ *=* .199, *CI* = .122-.519), following stimulation participants in the high expectation condition performed better than those in the low expectation condition (*t*_(86)_ = 2.294, *p* = .024; *M*_*diff*_ *=* .387, *CI* = .052-.722; Cohen’s *d* = .49). We did not find a main effect of expectation priming (*F*_[1,84]_ = 3.394, *p* = .069), of stimulation (*F*_[1,84]_ = .113, *p* = .737), interactions between time and stimulation (*F*_[1,84]_ = .191, *p* = .663) or expectation priming and stimulation (*F*_[1,84]_ = .620, *p* = .433), or a three-way interaction between time, priming, and stimulation (*F*_[1,84]_ = 1.057, *p* = .307).

### Does perceived application of non-invasive brain stimulation influence cognitive performance?

Following the stimulation procedure, we asked all participants if they believed their brain was genuinely stimulated during the course of the study. The majority of participants in each of the experimental conditions (67-81%) believed their brains were stimulated during the study; these proportions did not significantly differ between the experimental conditions. Moreover, repeating our analyses based on this perceived application of stimulation (i.e., “belief”) revealed no significant effect of belief on neuropsychological or WM performance. Similarly, repeated measures ANOVA examining expectation ratings in the different cognitive domains did not reveal significant differences at baseline, after receiving the priming message, or following stimulation based on belief.

### Do expectations or performance influence subjective experience?

In addition to asking about perceived nature of stimulation, we asked participants to rate their experience with the brain stimulation program. MANOVA across expectation priming and stimulation conditions revealed a significant effect of expectation priming on subjective experience (*F*_[7,78]_ = 2.263, *p* = .038, η_p_^2^ = .169).

After correcting for multiple comparisons, we found that participants who received high expectation priming reported higher levels of enjoyment (*F*_[1,84]_ = 9.737, *p* = .002, η_p_^2^ = .104), engagement (*F*_[1,84]_ = 7.183, *p* = .009, η_p_^2^ = .079), motivation (*F*_[1,84]_ = 8.146, *p* = .005, η_p_^2^ = .088), and satisfaction (*F*_[1,84]_ = 8.169, *p* = .005, η_p_^2^ = .089), compared to participants in the low expectation priming condition. Subjective experience did not differ based on stimulation condition, and did not significantly influence cognitive performance or expectation ratings at baseline. In an exploratory analysis of these feedback ratings, we did not find differences between participants who believed they received stimulation vs. those who did not.

### Are any individual factors associated with expectations or performance?

Based on our previous studies (Rabipour et al., 2017; Rabipour & Davidson, 2015), we examined whether prior knowledge of or experience with brain stimulation might influence expectations or cognitive performance. About one third of participants (30%) had previously heard of brain stimulation, and 20% had prior exposure to media reports about cognitive stimulation, but only 6% had any prior experience with brain stimulation. These proportions did not differ between experimental groups. Conversely, most participants (57%) reported having online gaming experience; the proportion was highest in people assigned to receive high expectation priming and active stimulation (21/24 = 88%; *X*^2^ = 12.924, *p* = .005). Neither expectations nor cognitive performance differed based on these individual factors.

We further explored whether gender might have an effect on our outcomes. Including gender as a variable in the repeated measures ANOVA for expectations ratings revealed no difference between women and men. However, adding gender as a variable in the ANOVA examining executive function composite scores across time revealed a significant main effect of gender (*F*_[1,80]_ = 4.143, *p* = .045, η_p_^2^ = .049) and significant three-way interaction between time, stimulation condition, and gender (*F*_[1,80]_ = 4.631, *p* = .034, η_p_^2^ = .055). Breaking down executive function scores across gender in each stimulation condition revealed a significant main effect of gender in participants who received sham stimulation (*F*_[1,39]_ = 7.714, *p* = .008, η_p_^2^ = .165), with higher scores in men both at baseline (*t*_(41)_ = 2.55, *p* = .015; *M*_*diff*_ = .51, *CI* = .105-.905; Cohen’s *d* = 0.78) and following stimulation (*t*_(41)_ = 2.797, *p* = .008; *M*_*diff*_ = .654, *CI* = .182-1.126; Cohen’s *d* = 0.84); we did not find an effect of gender in participants who received active tDCS (*F*_[1,41]_ = .123, *p* = .728). We did not find significant gender differences on n-back performance when adding gender as a factor to the ANOVA examining correct responses and false alarm rate across stimulation and priming conditions.

We did not find any group differences in reported levels of computer exposure, programming experience, or concern over declining cognitive functions (Table 3).

**Table 3.** Individual characteristics of participants across experimental conditions. Numbers represent the largest reported proportion per category.

|  | Computer Exposure: >2h per day | Programming Experience: None | Concern over Cognitive Functions: None |
| --- | --- | --- | --- |
| Active, High | 21/23 (91%) | 14/23 (61%) | 21/23 (91%) |
| Active, Low | 13/21 (62%) | 14/21 (67%) | 18/21 (86%) |
| Sham, High | 16/21 (76%) | 13/21 (62%) | 20/21 (95%) |
| Sham, Low | 17/23 (74%) | 16/23 (70%) | 20/23 (87%) |

### Did anticipated or experienced side effects interact with expectations or performance?

We asked participants about anticipated and experienced side effects before and after the stimulation procedure, respectively (Table 4). In each group, about half of participants or fewer (≤54%) anticipated experiencing side effects. After experiencing the stimulation procedure, however, the majority of participants in all groups (≥82%) – notably, all participants who received low expectation priming and active stimulation – reported experiencing at least one side effect. Similarly, the reported intensity was mild in the majority of participants in each group who reported experiencing side effects (≥67%), except for participants who received low expectation priming and active stimulation, who reported “considerable” intensity (35%), followed by “moderate” (29%), “mild” (24%) and “strong” (12%); this difference was significant compared to people who received high expectation priming and active stimulation (*X*^2^ = 8.316, *p* = 0.04). Moreover, we found a significant difference in reports of itchiness in people who received active stimulation, wherein a larger proportion of people who received low expectation priming reported experiencing itchiness, compared to those who received high expectation priming (*X*^2^ = 10.286, *p* = 0.001). Similarly, of people who received active stimulation, a larger proportion of people who received low (compared to high) expectation priming reported experiencing warmth or heat (*X*^2^ = 6.747, *p* = 0.009). The anticipation or overall experience of side effects did not significantly differ between groups, and did not significantly correlate with differences in expectation ratings, executive function scores, or WM performance.

**Table 4.** Proportion of participants who reported anticipating and experiencing side effects.

|  | **Anticipated Side Effects** | **Experienced: Itchiness** | **Experienced: Warmth / heat** | **Experienced: Pinching** | **Experienced: Pain** | **Experienced: Iron taste** | **Experienced: Fatigue** |
| --- | --- | --- | --- | --- | --- | --- | --- |
| Active, High | 12/23 (52%) | 8/22 (36%) | 7/22 (32%) | 5/22 (23%) | 2/22 (9%) | 0/22 (0%) | 2/22 (9%) |
| Active, Low | 7/21 (33%) | 17/20 (85%) | 15/21 (71%) | 7/21 (33%) | 7/21 (33%) | 1/21 (5%) | 4/21 (19%) |
| Sham, High | 5/20 (25%) | 13/20 (65%) | 8/20 (40%) | 3/20 (15%) | 2/20 (10%) | 1/20 (5%) | 2/20 (10%) |
| Sham, Low | 7/23 (30%) | 9/22 (41%) | 10/23 (44%) | 10/23 (44%) | 5/23 (22%) | 1/23 (4%) | 1/23 (4%) |

For clarity, we summarize our most pertinent findings in Table 5; see above for greater detail on each finding.

**Table 5.** Summary of results per experimental condition, with respect to: a) expectation ratings; b) composite Z scores on the executive function transfer tasks, with all scores normalized to baseline; c) proportion of correct trials (i.e., correct identification of targets and rejection of distractors) and false alarms (i.e., incorrect identification of distractors as targets, as a proportion of incorrect trials) on the WM task; and d) feedback ratings of subjective experience.

| **Main Finding** |  |
| --- | --- |
| **a) Expectations of Outcomes** | • Ratings largely "neutral" at baseline<br>• Ratings increase after high expectation priming and decrease after low expectation priming<br>• Following stimulation, ratings decrease across all groups |
| **b) N-back performance** | • Number of correct trials lowest in people who received active stimulation and low expectation priming<br>• Number of false alarms comparable across all groups |
| **c) Executive Function Scores** | • Performance similar between groups at baseline<br>• Following stimulation, improvement greater after receiving high (vs. low) expectation priming<br>• No difference in performance based on stimulation |
| **d) Subjective experience** | • Greater perceived enjoyment, engagement, motivation, and overall satisfaction in participants who received high (vs. low) expectation priming<br>• No group differences in perceived challenge, frustration, or boredom with the tasks |

## Discussion

Attitudes towards an intervention may greatly impact outcomes through placebo-like effects (Benedetti, Carlino, & Pollo, 2011). Factors such as expectation and satisfaction have potentially important implications for intervention adoption and adherence: for example, high expectations of positive outcomes may lead people to select and invest more time and resources in one program over another, and enhance their responsiveness to treatment. Conversely, low expectations may deter people from pursuing or adhering to an intervention. Given the current uncertainties surrounding tDCS mechanisms and the inconsistent findings in this field, understanding the potential interactions between expectations and stimulation is crucial.

Here we found that healthy subjects are generally uncertain about the effectiveness of NIBS in improving cognitive performance based on prior knowledge or familiarity with the concept. These neutral baseline ratings replicate our previous findings from people who completed our expectation scale as part of an online survey (Rabipour et al., 2017), suggesting that uncertainty about NIBS effectiveness extends to people who are willing to participate in such studies. Baseline ratings of perceived brain stimulation effectiveness did not appear to be influenced by exposure to advertisements and other media reports claiming that brain stimulation can improve cognitive function. Notably, very few participants reported having any prior experience with NIBS.

Our results further suggest that expectations of NIBS are malleable in response to simple written messages, even with a single exposure. Specifically, we demonstrate that reading information supporting the effectiveness of brain stimulation may induce higher expectations of improved cognitive performance, as well as improvements on objective measures of cognitive function. Conversely, reading information criticizing brain stimulation appears to decrease people’s expectations of brain stimulation effectiveness, and may subtly interfere with interventions aimed at improving cognitive performance. These patterns appear irrespective of whether the tDCS is genuinely applied throughout the session (i.e., active versus sham stimulation), and may be particularly pronounced in people who receive active stimulation.

Expectations of outcomes may influence cognitive performance. We found a subtle interaction between expectation priming and stimulation condition on both the neuropsychological transfer tests and the WM task performed concurrently with stimulation. Performance was greatest in people who received high, compared to low, expectation priming, irrespective of stimulation condition, suggesting a possible benefit of high expectations over stimulation. Conversely, participants randomly assigned to receive low expectation priming and active stimulation had the lowest performance scores on the WM task, suggesting that stimulation may have interfered with performance. Moreover, we found no evidence of improvement on the neuropsychological tests in participants assigned to receive low expectation priming, irrespective of active or sham stimulation. These effects are unlikely to result from knowledge of assignment to experimental condition (i.e., inadequate blinding), with the majority of participants believing that current was genuinely applied throughout the stimulation procedure.

Although differences in baseline performance on the neuropsychological tests were not statistically significant, a potential consideration is that participants randomly assigned to receive high expectation priming had slightly higher baseline scores than those assigned to the low expectation priming condition. Differences in baseline performance may explain why the interaction between time and expectation priming became marginal when examining performance on executive function transfer tests using a more conservative approach (i.e., with the condensed composite scores). Only future work will clarify this.

Notably, we found no evidence for an effect of stimulation condition on either the online WM task or the transfer measures of executive function. Coupled with the possible influence of expectations on outcomes, these findings put into the question the nature of reported stimulation effects in other studies examining tDCS and cognitive function.

Importantly, we found that people who received high expectation priming reported having a more positive experience, including greater enjoyment, engagement with the program, motivation to perform well, and satisfaction with the program. These findings have important potential implications for multi-session studies, which are becoming increasingly common: higher expectations of outcomes, either based on prior knowledge or delivered through researchers and clinicians – directly, via information provided, or indirectly, based on their attitudes and perceptions – may lead participants to be more enthusiastic after the first session and indirectly influence other outcomes.

Reports of anticipated and experienced side effects did not significantly differ across most conditions, although reports were greatest among participants assigned to receive low expectation priming and active stimulation. This may reflect a larger degree of surprise regarding the experience of sensations as well as greater allocation of attention to the sensations during task performance in these participants, who may have expected to receive sham stimulation based on their priming message. However, although side effects may have contributed to differences in performance and perceived experience, they are unlikely to have been the sole contributor to the group differences we found.

Despite the overarching uncertainty about NIBS outcomes at the outset of the tDCS procedure, and the malleability of expectations following expectation priming with the High and Low expectation messages, we found that all participants reported lower expectations of effectiveness following the stimulation procedure. This finding may reflect perceived difficulty of the WM task completed concurrently with tDCS, as the majority of participants expressed low confidence in their performance, as well as discomfort based on experienced side effects. However, factors such as perceived frustration and motivation did not account for differences in cognitive performance following tDCS in our sample. Taken together, our results suggest that expectations of NIBS may shape cognitive function independently of perceived performance.

### Limitations

Due to the subjective nature of surveys, we cannot rule out the possibility that our participants simply provided the responses they believed we wanted (i.e., “demand characteristics”). However, differences in cognitive performance outcomes following our balanced-placebo protocol offer evidence of a potential causal influence of expectations on cognitive performance following tDCS. Moreover, although our selected outcome measures (n-back task of WM and transfer tests of executive function) are commonly used in studies of cognitive function and NIBS, they may have been too coarse to detect subtle differences based on our experimental conditions. Our use of a single-task paradigm, rather than dual task, may explain why our results did not replicate those of Martin et al. (2014). Similarly, other methods of priming may be more powerful in creating an effect on stimulation outcomes. Finally, our sample of undergraduate students was relatively uniform, with few differences in background characteristics that may play a role in expectations of brain stimulation (e.g., education level, education type, culture, computer exposure, etc.). Thus, findings of this study are not necessarily representative of the general population. Related to the issue of sample composition is the issue of sample size. Although we determined sample size based on a priori power analyses, as with any study it is possible that a larger sample would have helped make our results easier to interpret. For example, in earlier analyses we included data on correct n-back trials from a participant who only had partial data available (i.e., no data on false alarm rate). The addition of this single data point made our results more pronounced, leading to a significant effect of expectation priming and larger interaction effect between expectation priming and stimulation condition on n-back performance. We ultimately decided to omit such data from our final analyses for a more conservative approach (see Materials and Methods), yielding the reported results. This example highlights issues regarding statistical power endemic to tDCS research.

### Significance for real-world trials

Our findings highlight the urgent need to consider the potential long-term effects of people’s beliefs in clinical trials and protocols. Here we were able to shape healthy subjects’ expectations with a simple, brief message, in just one lab session. These effects might be even greater in real-world situations, particularly for programs or treatments that require long-term compliance in patients seeking treatment for their condition. Trials may succeed or fail due to people’s pre-existing beliefs regarding the effectiveness of treatment received. In addition to a possible direct influence on outcomes, satisfaction with a program (or lack thereof) can potentially amplify results by positively or negatively affecting attrition rates and participant persistence over the course of treatment. Without evaluating the potential influence of such factors, the therapeutic viability of tDCS and optimal treatment parameters will remain unclear.

## Conclusions

The present study is the first, to our knowledge, to directly examine expectations of NIBS outcomes in the lab setting. Our overt attempt to prime expectations revealed subtle evidence for a potential causal influence of expectations on cognitive performance following tDCS. Notably, we found no group differences on the basis of stimulation. Results from our cognitive tasks suggest that positive priming may have beneficial effects on intervention experience and subsequent performance, whereas low expectations might be counterproductive in some cases (e.g., during administration of tDCS).

Replications of this study should examine the conditions under which expectations and stimulation meaningfully interact. Similarly, long-term investigations involving multiple sessions would further help elucidate the sustainability and potential additive effects of NIBS and expectations on cognitive performance. As such NIBS approaches become more common, people may form expectations that could interact with other intervention outcomes.

Here we show the importance of assessing participant expectations as a possible influence on the outcomes of cognitive interventions, and examining potential interactions between expectations and NIBS or related approaches to cognitive enhancement. Given the widespread appeal of NIBS as an affordable and accessible intervention in both healthy and clinical populations, understanding the potential impact of expectations is imperative to intervention design and for assessing the practical value of specific techniques.

## Acknowledgements

We thank Anita Popescu and Vilma Bezerra Alves for help with data processing, as well as Sima Sadeghinejad, Shreya Sreekantaswamy, and members of the Neuromodulation Lab at UCLA for help with data collection. We also thank Dr. Walter Dunn for helpful feedback on study design. For generous support the authors also wish to thank the Brain Mapping Medical Research Organization, Brain Mapping Support Foundation, Pierson-Lovelace Foundation, The Ahmanson Foundation, William M. and Linda R. Dietel Philanthropic Fund at the Northern Piedmont Community Foundation, Tamkin Foundation, Jennifer Jones-Simon Foundation, Capital Group Companies Charitable Foundation, Robson Family and Northstar Fund, as well as the Natural Sciences and Engineering Research Council (NSERC, Discovery Grant), Ontario Graduate Scholarships, and Fonds de Recherche Québec – Santé.

## Footnotes

1 Because we erred in not distinguishing between hits and correct rejections in programming our database, we were unable to analyse n-back performance as a function of hits vs. false alarms. We did, however, distinguish false alarms from misses in incorrect trials. Therefore, we were unable to perform sensitivity and bias analyses.

## References

Au, J., Katz, B., Buschkuehl, M., Bunarjo, K., Senger, T., Zabel, C., … Jonides, J. (2016). Enhancing Working Memory Training with Transcranial Direct Current Stimulation. Journal of Cognitive Neuroscience, 28(9), 1419-1432. doi: 10.1162/jocn_a_00979

Axelrod, B. N., Fichtenberg, N. L., Millis, S. R., & Wertheimer, J. C. (2006). Detecting incomplete effort with digit span from the Wechsler Adult Intelligence Scale - Third edition. Clinical Neuropsychologist, 20(3), 513-523. doi: Doi 10.1080/13854040590967117

Benedetti, F., Carlino, E., & Pollo, A. (2011). How Placebos Change the Patient’s Brain. Neuropsychopharmacology, 36(1), 339-354. doi: Doi 10.1038/Npp.2010.81

Bennabi, D., Pedron, S., Haffen, E., Monnin, J., Peterschmitt, Y., & Van Waes, V. (2014). Transcranial direct current stimulation for memory enhancement: from clinical research to animal models. Front Syst Neurosci, 8, 159. doi: 10.3389/fnsys.2014.00159

Berlim, M. T., Van den Eynde, F., & Daskalakis, Z. J. (2013). Clinical utility of transcranial direct current stimulation (tDCS) for treating major depression: A systematic review and meta-analysis of randomized, double-blind and sham-controlled trials. Journal of Psychiatric Research, 47(1), 1-7. doi: 10.1016/j.jpsychires.2012.09.025

Bikson, M., Grossman, P., Thomas, C., Zannou, A. L., Jiang, J., Adnan, T., … Woods, A. J. (2016). Safety of Transcranial Direct Current Stimulation: Evidence Based Update 2016. Brain Stimul, 9(5), 641-661. doi: 10.1016/j.brs.2016.06.004

Chew, T., Ho, K. A., & Loo, C. K. (2015). Inter- and Intra-individual Variability in Response to Transcranial Direct Current Stimulation (tDCS) at Varying Current Intensities. Brain Stimulation, 8(6), 1130-1137. doi: 10.1016/j.brs.2015.07.031

Coffman, B. A., Clark, V. P., & Parasuraman, R. (2014). Battery powered thought: Enhancement of attention, learning, and memory in healthy adults using transcranial direct current stimulation. Neuroimage, 85, 895-908. doi: 10.1016/j.neuroimage.2013.07.083

Crowe, S. F. (2000). Does the letter number sequencing task measure anything more than digit span? Assessment, 7(2), 113-117. doi: Doi 10.1177/107319110000700202

Dockery, C. A., Hueckel-Weng, R., Birbaumer, N., & Plewnia, C. (2009). Enhancement of Planning Ability by Transcranial Direct Current Stimulation. Journal of Neuroscience, 29(22), 7271-7277. doi: Doi 10.1523/Jneurosci.0065-09.2009

Farah, M. J. (2015). The unknowns of cognitive enhancement. Science, 350(6259), 379-380. doi: 10.1126/science.5893

Filmer, H. L., Dux, P. E., & Mattingley, J. B. (2014). Applications of transcranial direct current stimulation for understanding brain function. Trends Neurosci, 37(12), 742-753. doi: 10.1016/j.tins.2014.08.003

Foroughi, C. K., Monfort, S. S., Paczynski, M., McKnight, P. E., & Greenwood, P. M. (2016). Placebo effects in cognitive training. Proc Natl Acad Sci U S A, 113(27), 7470-7474. doi: 10.1073/pnas.1601243113

Freitas, C., Mondragon-Llorca, H., & Pascual-Leone, A. (2011). Noninvasive brain stimulation in Alzheimer’s disease: Systematic review and perspectives for the future. Experimental Gerontology, 46(8), 611-627. doi: 10.1016/j.exger.2011.04.001

Gandiga, P. C., Hummel, F. C., & Cohen, L. G. (2006). Transcranial DC stimulation (OCS): A tool for double-blind sham-controlled clinical studies in brain stimulation. Clinical Neurophysiology, 117(4), 845-850. doi: Doi 10.1016/J.Clinph.2005.12.003

Iyer, M. B., Mattu, U., Grafman, J., Lomarev, M., Sato, S., & Wassermann, E. M. (2005). Safety and cognitive effect of frontal DC brain polarization in healthy individuals. Neurology, 64(5), 872-875. doi: 10.1212/01.WNL.0000152986.07469.E9

Jones, K. T., Gozenman, F., & Berryhill, M. E. (2015). The strategy and motivational influences on the beneficial effect of neurostimulation: A tDCS and fNIRS study. Neuroimage, 105, 238-247. doi: 10.1016/j.neuroimage.2014.11.012

Kekic, M., Boysen, E., Campbell, I. C., & Schmidt, U. (2016). A systematic review of the clinical efficacy of transcranial direct current stimulation (tDCS) in psychiatric disorders. Journal of Psychiatric Research, 74, 70-86. doi: 10.1016/j.jpsychires.2015.12.018

Kincses, T. Z., Antal, A., Nitsche, M. A., Bartfai, O., & Paulus, W. (2004). Facilitation of probabilistic classification learning by transcranial direct current stimulation of the prefrontal cortex in the human. Neuropsychologia, 42(1), 113-117. doi: Doi 10.1016/S0028-3932(03)00124-6

Lanting, S., Haugrud, N., & Crossley, M. (2009). The effect of age and sex on clustering and switching during speeded verbal fluency tasks. Journal of the International Neuropsychological Society, 15(2), 196-204. doi: Doi 10.1017/S1355617709090237

Li, L. M., Uehara, K., & Hanakawa, T. (2015). The contribution of interindividual factors to variability of response in transcranial direct current stimulation studies. Frontiers in Cellular Neuroscience, 9. doi: ARTN 181 10.3389/fncel.2015.00181

Lopez-Alonso, V., Cheeran, B., Rio-Rodriguez, D., & Fernandez-del-Olmo, M. (2014). Inter-individual Variability in Response to Non-invasive Brain Stimulation Paradigms. Brain Stimulation, 7(3), 372-380. doi: 10.1016/j.brs.2014.02.004

Luber, B., & Lisanby, S. H. (2014). Enhancement of human cognitive performance using transcranial magnetic stimulation (TMS). Neuroimage, 85, 961-970. doi: 10.1016/j.neuroimage.2013.06.007

Martin, D. M., Liu, R., Alonzo, A., Green, M., & Loo, C. K. (2014). Use of transcranial direct current stimulation (tDCS) to enhance cognitive training: effect of timing of stimulation. Experimental Brain Research, 232(10), 3345-3351. doi: Doi 10.1007/S00221-014-4022- X

McCabe, D. P., Roediger, H. L., McDaniel, M. A., Balota, D. A., & Hambrick, D. Z. (2010). The Relationship Between Working Memory Capacity and Executive Functioning: Evidence for a Common Executive Attention Construct. Neuropsychology, 24(2), 222-243. doi: 10.1037/a0017619

McCambridge, J., Witton, J., & Elbourne, D. R. (2014). Systematic review of the Hawthorne effect: New concepts are needed to study research participation effects. Journal of Clinical Epidemiology, 67(3), 267-277. doi: Doi 10.1016/J.Jclinepi.2013.08.015

McIntire, L. K., McKinley, R. A., Goodyear, C., & Nelson, J. (2014). A Comparison of the Effects of Transcranial Direct Current Stimulation and Caffeine on Vigilance and Cognitive Performance During Extended Wakefulness. Brain Stimulation, 7(4), 499-507. doi: 10.1016/j.brs.2014.04.008

Medina, J., & Cason, S. (2017). No evidential value in samples of transcranial direct current stimulation (tDCS) studies of cognition and working memory in healthy populations. Cortex, 94, 131-141. doi: 10.1016/j.cortex.2017.06.021

Monti, A., Ferrucci, R., Fumagalli, M., Mameli, F., Cogiamanian, F., Ardolino, G., & Priori, A. (2013). Transcranial direct current stimulation (tDCS) and language. Journal of Neurology Neurosurgery and Psychiatry, 84(8), 832-842. doi: Doi 10.1136/Jnnp-2012-302825

Nitsche, M. A., Cohen, L. G., Wassermann, E. M., Priori, A., Lang, N., Antal, A., … Pascual-Leone, A. (2008). Transcranial direct current stimulation: State of the art 2008. Brain Stimul, 1(3), 206-223. doi: 10.1016/j.brs.2008.06.004

Penolazzi, B., Pastore, M., & Mondini, S. (2013). Electrode montage dependent effects of transcranial direct current stimulation on semantic fluency. Behavioural Brain Research, 248, 129-135. doi: 10.1016/j.bbr.2013.04.007

Poreisz, C., Boros, K., Antal, A., & Paulus, W. (2007). Safety aspects of transcranial direct current stimulation concerning healthy subjects and patients. Brain Res Bull, 72(4-6), 208-214. doi: 10.1016/j.brainresbull.2007.01.004

Rabipour, S., Andringa, R., Boot, W. R., & Davidson, P. S. R. (2017). What Do People Expect of Cognitive Enhancement? Journal of Cognitive Enhancement, 1-8. doi: 10.1007/s41465-017-0050-3

Rabipour, S., & Davidson, P. S. R. (2015). Do you believe in brain training? A questionnaire about expectations of computerised cognitive training. Behavioural Brain Research, 295, 64-70. doi: 10.1016/j.bbr.2015.01.002

Rabipour, S., Davidson, P. S. R., & Kristjansson, E. (2018). Measuring Expectations of Cognitive Enhancement: Item Response Analysis of the Expectation Assessment Scale. Journal of Cognitive Enhancement. doi: 10.1007/s41465-018-0073-4

Redick, T. S., & Lindsey, D. R. B. (2013). Complex span and n-back measures of working memory: A meta-analysis. Psychonomic Bulletin & Review, 20(6), 1102-1113. doi: 10.3758/s13423-013-0453-9

Riggall, K., Forlini, C., Carter, A., Hall, W., Weier, M., Partridge, B., & Meinzer, M. (2015). Researchers’ perspectives on scientific and ethical issues with transcranial direct current stimulation: An international survey. Scientific Reports, 5. doi: ARTN 10618 10.1038/srep10618

Ruf, S. P., Fallgatter, A. J., & Plewnia, C. (2017). Augmentation of working memory training by transcranial direct current stimulation (tDCS). Scientific Reports, 7. doi: ARTN 876 10.1038/s41598-017-01055-1

Sandrini, M., Brambilla, M., Manenti, R., Rosini, S., Cohen, L. G., & Cotelli, M. (2014). Noninvasive stimulation of prefrontal cortex strengthens existing episodic memories and reduces forgetting in the elderly. Frontiers in Aging Neuroscience, 6(289), 1-9. doi: 10.3389/fnagi.2014.00289

Sarkar, A., Dowker, A., & Cohen Kadosh, R. (2014). Cognitive enhancement or cognitive cost: trait-specific outcomes of brain stimulation in the case of mathematics anxiety. J Neurosci, 34(50), 16605-16610. doi: 10.1523/JNEUROSCI.3129-14.2014

Scheldrup, M., Greenwood, P. M., McKendrick, R., Strohl, J., Bikson, M., Alam, M., … Parasuraman, R. (2014). Transcranial direct current stimulation facilitates cognitive multi-task performance differentially depending on anode location and subtask. Front Hum Neurosci, 8, 665. doi: 10.3389/fnhum.2014.00665

Schwarz, K. A., Pfister, R., & Buchel, C. (2016). Rethinking Explicit Expectations: Connecting Placebos, Social Cognition, and Contextual Perception. Trends in Cognitive Sciences, 20(6), 469-480. doi: 10.1016/j.tics.2016.04.001

Shiozawa, P., Duailibi, M. S., da Silva, M. E., & Cordeiro, Q. (2014). Trigeminal nerve stimulation (TNS) protocol for treating major depression: An open-label proof-of-concept trial. Epilepsy Behav, 39, 6-9. doi: 10.1016/j.yebeh.2014.07.021

St Clair-Thompson, H. L. (2010). Backwards digit recall: A measure of short-term memory or working memory? European Journal of Cognitive Psychology, 22(2). doi: 10.1080/09541440902771299

Underwood, E. (2016). Cadaver study casts doubts on how zapping brain may boost mood, relieve pain. Science. Retrieved from doi: 10.1126/science.aaf9938

Voroslakos, M., Takeuchi, Y., Brinyiczki, K., Zombori, T., Oliva, A., Fernandez-Ruiz, A., … Berenyi, A. (2018). Direct effects of transcranial electric stimulation on brain circuits in rats and humans. Nat Commun, 9(1), 483. doi: 10.1038/s41467-018-02928-3

Walsh, V. Q. (2013). Ethics and Social Risks in Brain Stimulation. Brain Stimulation, 6(5), 715- 717. doi: 10.1016/J.Brs.2013.08.001

Zhao, H. C., Qiao, L., Fan, D. Q., Zhang, S. Y., Turel, O., Li, Y. H., … He, Q. H. (2017). Modulation of Brain Activity with Noninvasive Transcranial Direct Current Stimulation (tDCS): Clinical Applications and Safety Concerns. Frontiers in Psychology, 8. doi: 10.3389/fpsyg.2017.00685

